# Expression and role of the RNA-binding protein, ZFP36L1, in mouse T follicular helper cell differentiation and function

**DOI:** 10.64898/2026.02.02.703435

**Authors:** Helena A. Carslaw, Sam Woolliscroft, Emily M. Watson, Sarah E. Bell, Michelle A. Linterman, Martin Turner, Louise M. C. Webb

## Abstract

T follicular helper (Tfh) cells are critical for germinal centers (GC), the specialised microenvironment where long-lived humoral immunity is generated in response to vaccination or infection. Within the GC, B cells engage with Tfh cells and elicit their help in the form of cytokines and cell-surface co-stimulator molecules. Tfh helper activity must be rapidly available in response to B cell engagement, yet the mechanisms controlling this helper activity remain poorly characterized. Post-transcriptional regulation of mRNA decay and translation offer one way to rapidly and temporally tune Tfh cell activity. ZFP36L1, a member of the ZFP family of RNA-binding proteins, is a candidate modulator of Tfh cell helper activity as it controls cytokine production and responses in other T cell lineages, modulating their differentiation and function. We sought to determine if ZFP36L1 is also important for Tfh cell biology. In this study, we show expression of ZFP36L1 by Tfh cells. We selectively delete ZFP36L1 from Tfh cells and analyze the effect this has on the GCs. Surprisingly, we find the GC response and affinity maturation is resilient to deletion of ZFP36L1 from Tfh cells.

## Introduction

Germinal centers (GCs) form in response to vaccination or infection. They are pivotal for the generation of robust and effective immune responses, generating life-long immunity via formation of memory B cells and antibody secreting cells (ASCs) (1). T follicular helper (Tfh) cells are a subset of CD4^+^ T cells essential for the development and function of GCs (2). Tfh cells interact with antigen-specific B cells both outside the GC at the T:B cell border, and within the light zone of the GC where they provide key helper signals via CD40 engagement and provision of cytokines. Rapid elicitation of Tfh cell help by engaged B cells is paramount for B cell entry into the GC and their subsequent rapid proliferation within it (1–3). However, this help must be carefully controlled to prevent aberrant immune responses and maintain the selective environment critical for ensuring affinity maturation of the B cell response (1) .

RNA binding proteins (RBPs) are candidate regulators of Tfh cell activity as they can provide the rapid response to stimuli required for T-B cell interactions within the GC. They orchestrate rapid cellular responses by eliciting post-transcriptional control of cytokines and other immune mediators (4). By controlling RNA translation and/or mediating its decay, RBPs play critical roles in lymphocyte biology both in terms of development and function (5). Thousands of RBPs have been annotated in T cells (6–8), over a hundred of which have been shown to bind to the untranslated region (UTR) of IFNγ, IL-2, and IL-10 mRNAs in human T cells suggesting the potential for exquisite time/stimuli-dependent control of protein production (10).

Amongst the RBP that interact with cytokine mRNAs is the Zinc Finger Protein (ZFP)36 family, which is comprised of ZFP36, ZFP36L1 and ZFP36L2. They are recognised as important for lymphocyte function, mediating their effects via regulation of cytokine mRNAs and other transcripts encoding transcription factors, cell cycle and metabolic regulators, where they play complex and overlapping roles(5). Whilst some responses of lymphocytes show unique dependencies on individual ZFP family members, others show redundancy (9–11). For example, deletion of ZFP36L1 from human T cells results in increased half-life of cytokine mRNA and this is attributed to its ability to destabilize mRNA (12), whilst in memory T cells ZFP36L2 suppresses the translation of IFNγ mRNA (13). However, deletion of both ZFP36L1 and ZFP36L2 in CD8^+^ T cells results in increased IFNγ production that is not seen with deletion of either ZFP family member alone (14). In CD8+ cells and T regulatory (Treg) cells, ZFP36L1 and ZFP36L2 are important for controlling responses to IL-2(15,16). In B cells, ZFP36L1 maintains the marginal zone B cell phenotype (17) and regulates plasma cell homing (18) but is dispensable for the GC reaction due to redundancy with ZFP36L2 (19). Although the ZFP family participates in a multitude of lymphocyte functions, their role in Tfh cells remains unknown.

Tfh cells share transcriptional signatures with other T-helper subsets and until recently, it was not possible to specifically manipulate these cells *in vivo* via cre-directed targeting of genes. We recently repurposed an *Il21cre* fate-mapping mouse to specifically target precursor-Tfh, Tfh, and GC-Tfh cells(20). Here, we use this *Il21cre* fate-mapping system to specifically delete *Zfp36l1* from Tfh cells allowing us to assess its role in Tfh cell development and function. We show that despite expression of ZFP36L1 in Tfh cells, its loss has no effect on GC formation or output, suggesting there may be redundancy in the ZFP family for Tfh cell function.

## Results and Discussion

### ZFP36L1 expression during Tfh cell differentiation

Differentiation of naïve CD4^+^ T cells into Tfh cells occurs via a multistep process dependent on cognate interactions with antigen presenting cells (APCs)(21). Naïve CD4^+^ T cells are first stimulated by APCs resulting in their activation and formation of precursor Tfh cells (pre-Tfh). Expression of CXCR5 and EBI2 by pre-Tfh cells enable their migration and relocation to the interfollicular area between the T and B cell zones of the draining lymph node (LN; (21)). CD4^+^ T cells have a high degree of plasticity and pass through a bipotent Th1/Tfh precursor state with subsequent engagement with B cells driving Tfh cell differentiation (22). A subset of Tfh cells then enter the GC (GC-Tfh) where they directly interact with GC B cells(23), driving the proliferation and affinity maturation of GC B cells (1–3). We first sought to examine expression of the *Zfp36, Zfp36l1, and Zfp36l2* transcripts during these distinct phases of Tfh cell differentiation. Using published data from a model of Tfh and Th1 cell bifurcation following *Plasmodium chabaudi* infection(24), we examined pseudo time analysis of *Zfp36, Zfp36l1, and Zfp36l2* expression by antigen-specific TCR transgenic PbTII CD4^+^ cells. This analysis showed that Tfh cells express *Zfp36, Zfp36l1, and Zfp36l2* mRNA and that expression is retained after the Th1/Tfh bifurcation point. Interestingly, *Zfp36l1* transcript expression closely mirrored that of the Tfh-defining transcription factor, *Bcl6* (Fig. 1A) suggesting a role for ZFP36L1 in Tfh differentiation/function. To determine if ZFP36L1 protein expression also accumulated in Tfh cells we used intracellular staining to of CD4^+^ T cells following immunisation with a T-dependent antigen, NP-KLH. Using well defined markers for non-Tfh, pre-Tfh, Tfh, GC-Tfh (GC-resident Tfh), and T follicular regulatory (Tfr) cells (20) we determined ZFP36L1 expression during different stages of Tfh cell differentiation. Lymphocytes were harvested from the inguinal (draining) LN (iLN) 10 days after subcutaneous immunisation with NP-KLH/Alhydrogel (a timepoint where all Tfh cell subsets would be present) and ZFP36L1 protein expression was determined (Fig. 1B). Non-Tfh, pre-Tfh, Tfh, GC-Tfh, and Tfr cells were gated as shown (Fig. 1C) and both the percentage of cells expressing ZFP36L1 and the levels of ZFP36L1 expression (MFI) determined (Fig.1D-F). ZFP36L1 expression was seen in >90% of non-Tfh, pre-Tfh, Tfh, GC-Tfh, and Tfr cells (Fig. 1D-F). Although there were only incremental increases in the proportion of ZFP36L1^+^ cells in each subset during Tfh differentiation (Fig. 1F), the amount of ZFP36L1 peaked at the Tfh stage of differentiation (Fig.1E) with an MFI nearly 2-fold greater than that of pre-Tfh, and non-Tfh cells. Thus, the majority of activated CD4^+^ T cells express ZFP36L1 protein following immunisation and ZFP36L1 expression is retained throughout the stages of Tfh cell differentiation, peaking prior to their entry into the GC. This suggests ZFP36L1 could play a role in the development and/or biology of the GC response.

**Figure 1.**
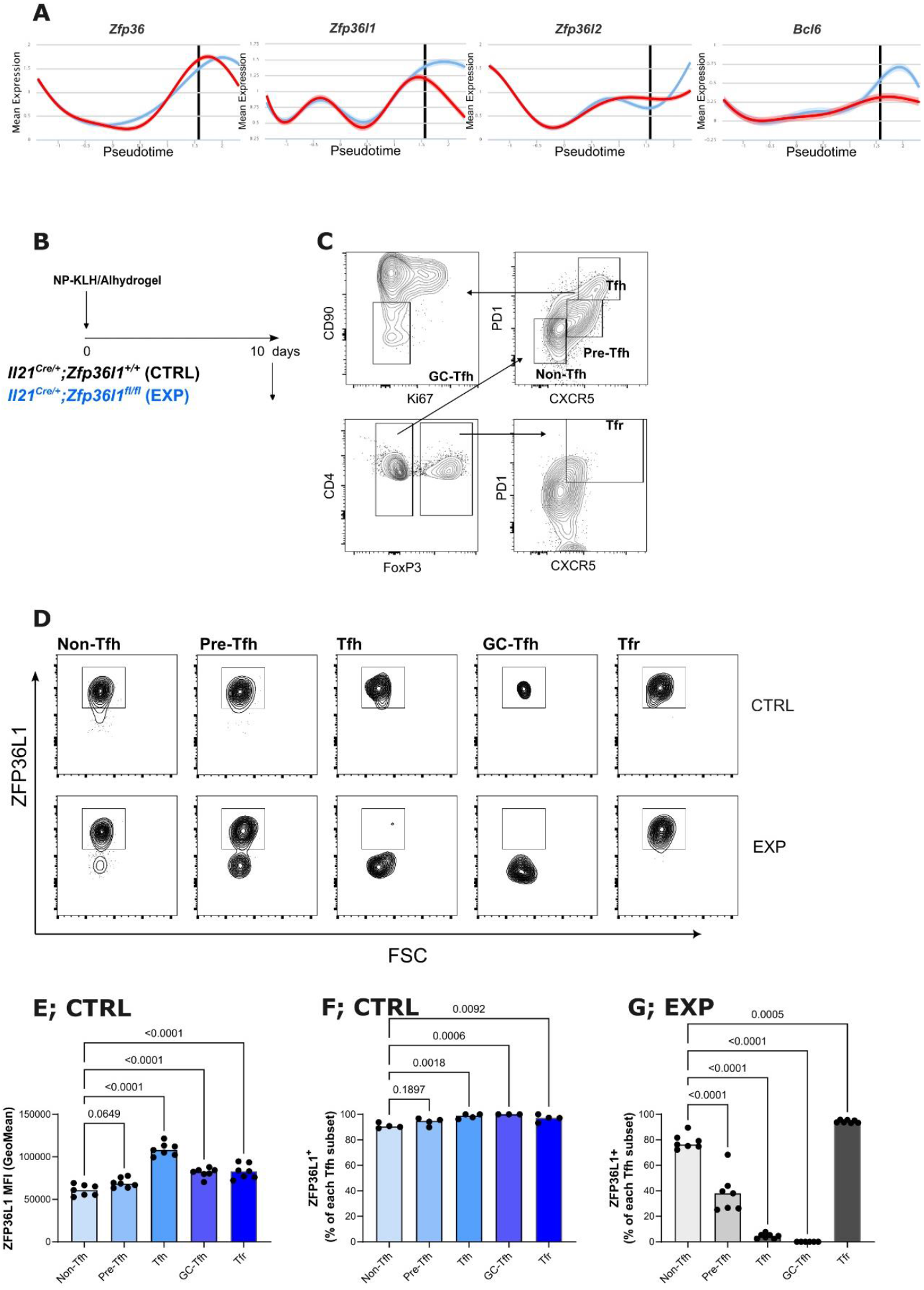
ZFP36L1 expression during Tfh cell differentiation. (**A**), Pseudo time analysis *Zfp36, Zfp36l1, Zfp36l2, and Bcl6* expression by antigen-specific TCR transgenic PbTII CD4^+^ cells following *P. chaubaudi* infection of mice. Relative expression levels of each gene in a single Tfh (blue line) or Th1 (red line) cells relative to pseudotime. (**B**), Experimental scheme for (**C-G**). *Il21*^*+/Cre*^*;Zfp36l1*^*+/+*^ (CTRL) and *Il21*^*+/Cre*^*;Zfp36l1*^*fl/fl*^ (EXP) mice were immunized subcutaneously with NP-KLH/Alhydrogel and draining inguinal lymph nodes (iLNs) harvested 10 days later. (**C**), Gating strategy used for (**E-G**). (**D**), Representative flow cytometry plots showing staining for ZFP36L1 in different Tfh subsets following 3 hour restimulation *ex vivo* for EXP or CTRL mice. (**E**) Bar graphs showing expression levels of ZFP36L1 in different Tfh subsets as determined by flow cytometry for CTRL mice. (**F, G**,) Bar graphs showing the percentage of cells expressing ZFP36L1 in CTRL (**F**) and EXP (**G**) mice. Bar graphs show medians with each symbol representing an individual mouse. Data is representative of 2 independent experiments with 4-8 mice per group.

We examined ZFP36L1 expression in a strain of mice designed to delete *Zfp36l1* in Tfh cells (Fig. 1, D, F). *Il21*-fate mapped (*Il21*-fm) mice enable Cre expression in Tfh cells, GC-Tfh cells and a proportion of pre-Tfh cells (20). Importantly, *Il21*-fate mapping spares suppressive Tfr cells, which function to balance Tfh cell helper function. In *Il21*^*+/Cre*^*;Zfp36l1*^*fl/fl*^ mice, we observed loss of ZFP36L1 protein from Tfh cells, GC-Tfh cells and >50% of pre-Tfh cells but it remained present in Tfr cells (Fig. 1E&H). The *Il21*^*+/Cre*^*;Zfp36l1*^*fl/fl*^ mice thereby provided a valuable tool to investigate the function of ZFP36L1 in Tfh cell differentiation and biology.

### Tfh differentiation and function in the absence of ZFP36L1

Using *Il21*-fm mice to specifically delete *Zfp36l1* from Tfh, GC-Tfh cells and a proportion of pre-Tfh cells, we first analysed Tfh cell differentiation by flow cytometry. Control (CTRL; *Il21*^*+/Cre*^*;Zfp36l1*^*+/+*^) or experimental (EXP; *Il21*^*+/Cre*^*;Zfp36l1*^*fl/fl*^) mice were immunised on both hind flanks with NP-KLH/Alhydrogel and iLNs harvested for flow cytometry and imaging 9 days later during the peak of the GC response (Fig. 2A). Flow cytometry analysis showed normal Tfh and Tfr cell differentiation (Fig.2B-C) with no perturbations in Tfh or GC-Tfh cells despite lacking ZFP36L1 expression. Enumeration of the different Tfh cell subsets (non-Tfh, pre-Tfh, Tfh, GC-Tfh, Tfr) confirmed this (Fig. 2C). In addition, there was no difference observed in CXCR3^+^ Th1 cells. Together, this shows that loss of ZFP36L1 does not affect Tfh cell differentiation or Th1/Tfh bifurcation (Fig. 2B,C).

**Figure 2.**
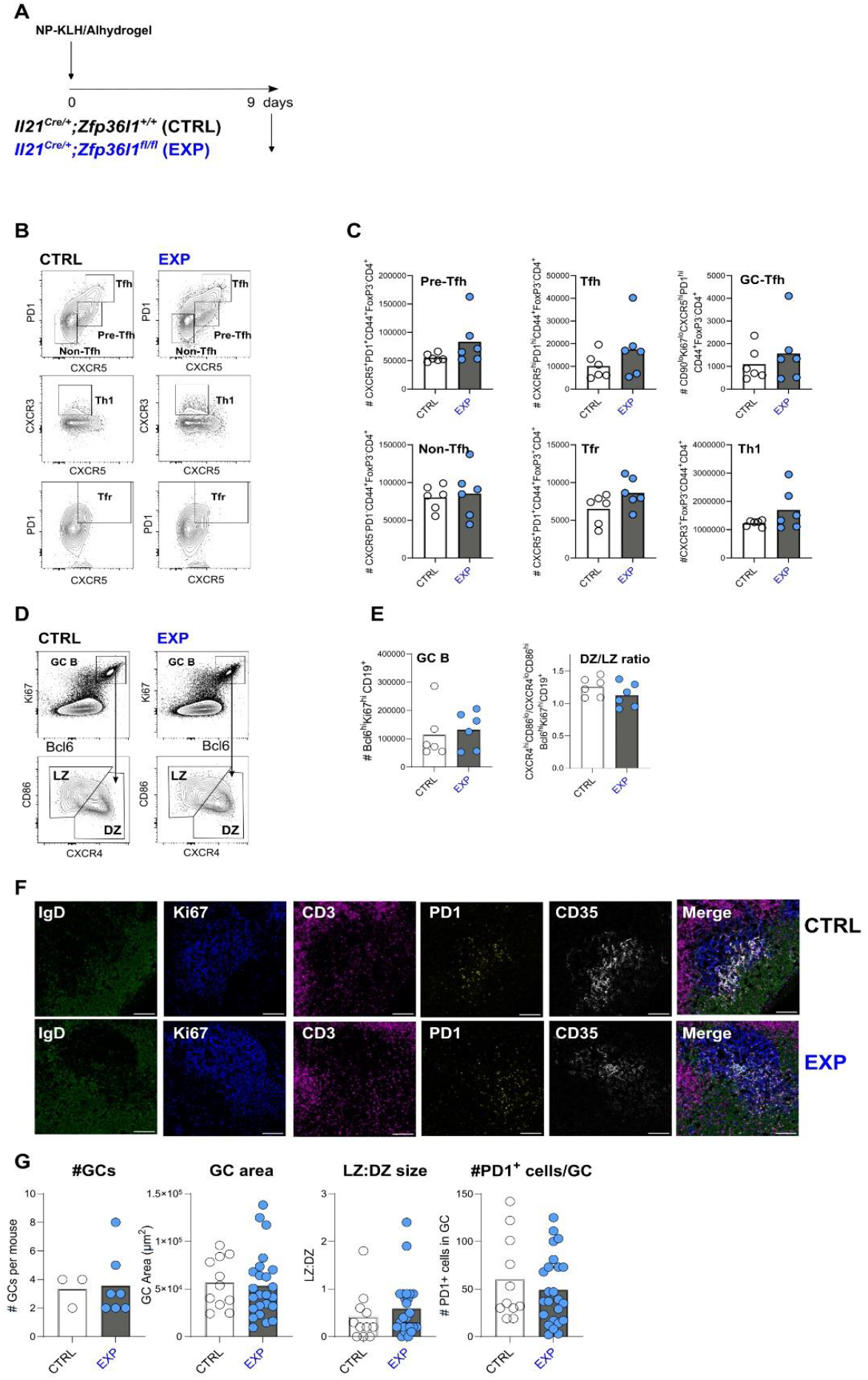
Tfh differentiation and function in the absence of ZFP36L1. (**A**), Experimental scheme for (**B-E**). *Il21*^*+/Cre*^*;Zfp36l1*^*+/+*^ (CTRL) and *Il21*^*+/Cre*^*;Zfp36l1*^*fl/fl*^ (EXP) mice were immunized subcutaneously with NP-KLH/Alhydrogel and draining inguinal lymph nodes (iLNs) harvested 9 days later. (**B**), Representative flow cytometry plots showing Tfh subsets, Th1 cells and Tfr cells in draining iLNs 9 days post immunisation (d.p.i.). (**C**), Bar graphs showing the number of different T cell populations in CTRL (white circles) and EXP (blue circles) mice. (**D**), Representative flow cytometry plots showing germinal center (GC) B cells and light zone (LZ) and dark zone (DZ) GC B cells for CTRL and EXP mice. (**E**), Bar graphs showing the number GC B cells and the ratio of DZ/LZ GC B cells in draining iLNs 9 d.p.i. in CTRL (white circles) and EXP (blue circles) mice. Data is representative of 2 independent experiments with 4-8 mice per group. (**F**), Representative confocal images of GCs from CTRL and EXP mice 9 d.p.i. IgD =green Ki67 = blue, CD3 = magenta, CD35 = white, PD1 = yellow). Scale bar = 100mm. GC area is defined as the combined area of Ki67+ area and CD35+ area. (**G**), Bar graphs showing the quantification of GC number, size, and number of PD1+ cells in GC from confocal images in CTRL (white circles) and EXP (blue circles) mice. Bar graphs show median values with each symbol representing an individual mouse (for # GCs/mouse) or each symbol represents a GC.

Since the main function of Tfh cells is to provide help to B cells responding to antigen, it was important to determine if ZFP36L1 played a role in Tfh cell function. In T-dependent responses, Tfh cells are critical for formation and maintenance of the GC, mainly due to their provision of helper signals that drive GC B cell proliferation (2,3). To assess the role of ZFP36L1 in Tfh cell activity, we first examined GC formation in *Il21*^*+/Cre*^*;Zfp36l1*^*fl/fl*^ (EXP) mice, comparing their responses to littermate controls (*Il21*^*+/Cre*^*;Zfp361*^*+/+*^; CTRL). As shown, GC B cells formed in *Il21*^*+/Cre*^*;Zfp36l1*^*fl/fl*^ mice (Fig. 2D) and with no changes in GC B cell numbers in mice lacking ZFP36L1 expression in their Tfh cells (Fig. 2E). Following reciept of helper signals from Tfh cells, GC B cells move to the dark zone (DZ) of the GC where very rapid diviision occurs (1). We subdivided GC B cells into light zone (LZ) or DZ phenotypes by flow cytometry but found no difference in the proportions of LZ and DZ GC B cells in *Il21*^*+/Cre*^*;Zfp36l1*^*fl/fl*^ mice compared to control mice (Fig. 2D-E). This suggests that in the absence of ZFP36L1, Tfh cells retain their ability to drive GC B cell proliferation. We confirmed these finding with confocal imaging analysis of iLNs where normal numbers and sizes of GCs were observed in mice with ZFP36L1-deficient Tfh cells (Fig. 2F-G).Additionally, the LZ:DZ ratio and the numbers of Tfh in each GC were also unaffected in *Il21*^*+/Cre*^*;Zfp36l1*^*fl/fl*^ mice compared to control mice (Fig. 2F-G).

### Affinity maturation is resilient to loss of ZFP36L1 expression in Tfh cells

The hallmark of the GC response is affinity maturation of the antibody response(25). This is dependent on Tfh cells providing helper signals to GC B cells in a competitive manner. It enables the evolution of the GC B cell response and selection of GC B cells with B cell receptors that bind the immunising antigen effectively enough to present it to Tfh cells (1). To test how ZFP36L1 .expression in Tfh cells contributes to affinity maturation, we measured the antibody response to NP-KLH over a 3-week period, allowing the evolution of the response to be characterised using ELISAs (for serum antibodies) and ELISpots (for antibody secreting cells) to measure total and high affinity antibodies to NP. Control (*Il21*^*+/Cre*^*;Zfp36l1*^*+/+*^) and experimental (*Il21*^*+/cre*^*;Zfp36l1*^*fl/fl*^) mice were immunised with NP-KLH/Alhydrogel on both flanks and blood was taken on days 7, 14, and 21 post immunisation (p.i.) (Fig. 3A). In addition, bone marrow antibody secreting cells (ASCs) with total and high affinity NP binding capabilities were measured by ELISpot at day 21 p.i. The levels of total NP20-binding antibodies and numbers of ASCs were not different between *Il21*^*+/Cre*^*;Zfp36l1*^*fl/fl*^ and control mice (Fig. 3B). Loss of ZFP36L1 in Tfh cells also had no impact on the generation of high affinity (NP2) binding antibodies (Fig. 3C). We determined the degree of affinity maturation occurring within individual mice by calculating the ratio of NP2/NP20 titers. This showed that the lack of ZFP36L1 in Tfh cells had no detectable effect on affinity maturation (Fig. 3D).

**Figure 3.**
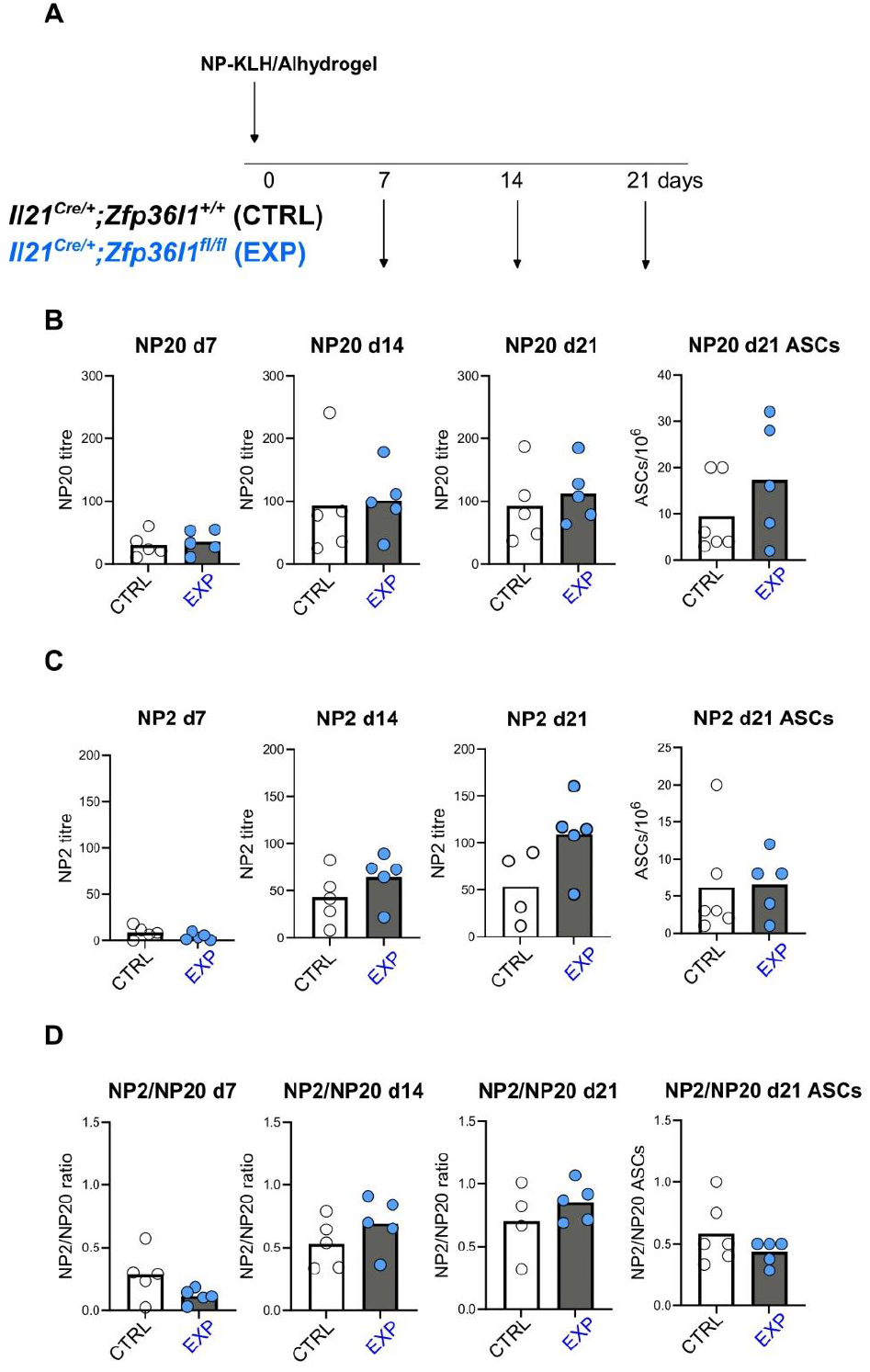
Affinity maturation is resilient to loss of ZFP36L1 expression in Tfh cells. (**A**), Experimental scheme for (**B-D**). *Il21*^*+/Cre*^*;Zfp36l1*^*+/+*^ (CTRL) and *Il21*^*+/Cre*^*;Zfp36l1*^*fl/fl*^ (EXP) mice were immunized subcutaneously with NP-KLH/Alhydrogel and blood taken on days 7,14, and 21 post immunisation (p.i.). with bone marrow harvested on day 21 p.i.. (**B**), Bar graphs showing titers of anti-NP20 IgG1 (total affinity) on day 7, 14 and 21 p.i and number of ASCs on day 21 p.i. from CTRL (white circles) and EXP (blue circles) mice. (**C**), Bar graphs showing titers of anti-NP2 IgG1 (high affinity) on day 7, 14 and 21 p.i and ASCs on day 21 p.i. from CTRL (white circles) and EXP (blue circles) mice. (**D**), Bar graphs showing affinity maturation as determined by titers of NP2 relative to NP20 antibodies and numbers of bone marrow ASCs from CTRL (white circles) and EXP (blue circles) mice. Bar graphs show median values with each symbol representing an individual mouse. Data is representative of 2 independent experiments with 4-8 mice per group.

## Concluding Remarks

The provision of rapid but controlled help to B cells during the GC response is paramount to an effective immune response. It facilitates the generation of B cells capable of producing antibodies that bind with sufficient affinity, avidity and selectivity for the immunizing antigen (20). Tfh and B cells engagement in the GC occurs by entanglement, resulting in rapid surface display of CD40L (and presumably other helper factors) by Tfh cells (26). Post-transcriptional regulation of Tfh helper factors offers a mechanism to ensure this rapid but temporal response. By using a model system where we could specifically ablate ZFP36L1 in Tfh cells, we were able to determine the role of this RBP in Tfh cell function. However, despite, expression of ZFP36L1 protein in Tfh cells, and evidence that *Zfp36l1* mRNA preferentially associates with Tfh cells after the Tfh/Th1 bifurcation point, there was no observable phenotype in *Il21*^*+/Cre*^*;Zfp36l1*^*fl/fl*^ mice in terms of Tfh cell differentiation, GC responses and affinity maturation.

It is probable that other ZFP family members compensate for a lack of ZFP36L1 in Tfh cells, as has been seen in other T cell subsets. As described above, deletion of both ZFP36L1 and ZFP36L2 results in far greater production of IFNγ by CD8^+^ cells than observed by deletion of just a single ZFP family member. Likewise, ZFP36L1 restrains CD8^+^ T cell cytotoxic function but it is only upon deletion of both ZFP36L1 and ZFP36L2 that this is revealed (10). Also, mice with Treg cells deficient in ZFP36L1 show a milder phenotype than that of mice with a combined deletion of ZFP36L1 and ZFP36L2 (15).

The mechanisms controlling elicitation of Tfh cell help are likely to be multilayered, acting both pre and post translationally on helper proteins. RNA-binding proteins, such as the ZFP family, potentially provide a key layer of regulation and remain promising candidates for controlling the elicitation of Tfh cell help for the GC B cell. The complex redundancy that exists within the ZFP family of RBPs suggests that their function in Tfh cells will most likely be revealed by deleting multiple family members.

## Data Limitations and Perspectives

Here, we focused on the effect of deleting the RNA-binding protein, ZFP36L1, from Tfh cells on a primary T cell dependent immune response. We compared the GC response mounted in ZFP36L1-sufficient and deficient Tfh cells using *Il21*-fatemapping to mark Tfh cells and their precursor populations. The lack of a measurable effect of ZFP36L1 deficiency on the GC response and affinity maturation may be masked by compensatory mechanisms that exist within the system. ZFP36L1 is a member of a family of RNA-binding proteins which may provide compensatory role.

## Materials and Methods

### Mice

*II21*^*+/Cre*^*;Zfp36l1*^*fl/fl*^ mice were bred and housed at the Babraham Institute Biological Support Unit. All mice were housed in individually ventilated cages, at an ambient temperature of 19-21^0^C, 52% relative humidity and under pathogen free conditions. All mice used in experiments were between 8-12 weeks of age and a mix of sexes. All mouse work and experimentation were approved by the Babraham Institute Animal Welfare and Ethical Review Body and complied with all European Union and United Kingdom Home Office legislation and standards (PPL: P4D4AF812 & PP9973990).

### Immunisations

Mice were immunised by subcutaneous injection of 50mg NP-KLH (N-5060-25, Biosearch Technologies) in a 1:1 ratio with 2% Alhydrogel (vac-alu-10, InvivoGen), 100ml per flank.

### Flow Cytometry

Cells were processed as previously described (20). In brief, inguinal lymph nodes (iLNs) were pushed through a 30mm cell strainer (1050552, CellTrics) to create a single cell suspension in FACS buffer (2% FBS (F9665, Sigma), in PBS with 0.05mM EDTA (15575020, Invitrogen)). Cells were counted with CASY TT Cell Counter (OLS OMNI Life Science) and plated in v-bottom 96-well plate (611V96, Thermo Fisher Scientific) at a concentration of 2x10^6^ cells/ml. Cells were stained with surface antibodies for 2 hours at 4^0^C, fixed with 2% formaldehyde (1.00496.0700, Sigma Aldrich) for 23 minutes at room temperature (RT), permeabilised with FoxP3 Permeabilization Buffer (00-8333-56, Invitrogen) and stained with intracellular antibodies overnight at 4^0^C. Antibodies are listed in Supp. Tables 1 and 2. For staining of ZFP36L1, cells were stimulated with PMA/Ionomycin (00-4970-93, Invitrogen) in complete RPMI (21875-034, Gibco) supplemented with 10% (v/v) FBS (F9665, Sigma), 1% (v/v) Penicillin-Streptomycin (15140-122, Thermo Fisher Scientific) and 55 µM β-mercaptoethanol (21985023, Thermo Fisher Scientific) for 3 hours at 37^0^C. Cells were stained and fixed as above but with surface antibody incubation time reduced to 45 minutes.

### Microscopy

iLNs were fixed with 4% PFA (J19943.K2, Thermo Fisher Scientific) for 2 hours at 4^0^C, washed briefly with PBS and left overnight in 30% sucrose (w/v) (S/8600/60, Fisher Chemical) to cryopreserve. iLNs were embedded in O.C.T medium (AGR1180, Agar Scientific), frozen on dry ice and sectioned at 10<u>µ</u>m at -20^0^C with a Leica Cryostat (Leica CM3050S). For immunofluorescence staining, slides were dried under airflow for 10 minutes, rehydrated in PBS-Tw20 (0.5% Tween-20 (P1379, Sigma Aldrich) (v/v) in PBS) for 15 minutes and blocked with 10% normal donkey serum (D9663, Sigma-Aldrich) (v/v) in 2% BSA (A7906, Sigma Aldrich in PBS for 45 minutes at RT. Slides were incubated with the primary antibodies listed in Supp. Table 3 for 1 hour at room temperature and secondary antibody listed in Supp. Table 3, for 30 minutes at room temperature. All antibodies were diluted in 1% BSA (w/v) in PBS-Tw20. Slides were washed with PBS-Tw20 for 5 minutes, 3 times between all staining and blocking steps. Finally, slides were washed briefly twice with PBS-Tw20 and twice with PBS before mounting with Hydromount mounting medium (S6-0106, Geneflow). Images were acquired on Leica Stellaris 8 confocal microscope using 20x/0.75 numerical aperture air lens. Every germinal center (GC) per LN was imaged at 512x512 resolution. Images were processed with ImageJ (v.2.14.0/1.54f) and quantitative analysis performed using a custom CellProfiler (v.4.2.4) pipeline. GCs were identified as CD35^+^ and/or Ki67^+^. Within the GC, the light zone was identified as CD35^+^ and dark zone as CD35^-^Ki67^+^. Tfh cells were identified as PD1^+^ and their numbers quantified within each GC compartment using CellProfiler.

### ELISAs

Flat-bottom Nunc Maxisorb plates (456537, Thermo Fisher Scientific) were coated with 50ml of either NP-20-BSA (10mg/ml, N5050G-100, Biosearch Technologies) or NP-2-BSA (2mg/ml, N5050L-100, Biosearch Technologies) diluted in PBS, overnight at 4^0^C. Plates were blocked with 200ml of 2% BSA (A7906, Sigma Aldrich, ) (w/v) in PBS-Tw20 (0.05% Tween-20 (P1379, Sigma Aldrich) (v/v) in PBS) for 2 hours at RT. Serum was serially diluted down the plate 1:3 from a starting concentration of 1:50 and incubated for 1 hour at RT. For detection of serum antibodies, the plate was incubated with 50ml anti-mouse IgG1-HRP (1/8000, ab97240, Abcam) for 1 hour at RT followed by 50ml TMB substrate (421101, BioLegend) until an appropriate dynamic range was reached. The reaction was quenched by addition of 50ml H_2_SO_4_ (2.5N) and optical density measurements were taken at 450nm on PHERAstar FS plate reader (BMG Labtech).

### ELISpot

96-well multiscreen HTS HA filter plates (MSHAS4510, Sigma Aldrich,) were coated with 50ml NP-2-BSA (2mg/ml, N5050L-100, Biosearch Technologies) or NP-20-BSA (10mg/ml, N5050G-100, Biosearch Technologies) overnight at 40C. Plates were then incubated with 50ml of complete RPMI (21875-034, Gibco) supplemented with 10% (v/v) FBS (F9665, Sigma), 1% (v/v) Penicillin-Streptomycin (15140-122, Thermo Fisher Scientific) and 55 µM β-mercaptoethanol (21985023, Thermo Fisher Scientific) for 1 hour at RT. Bone marrow was extracted from the hind legs of the mice and red blood cells were lysed by incubation with Ammonium-Chloride red blood cell lysis buffer for 5 minutes. Bone marrow cells were resuspended in complete RPMI medium and 2x106 cells plated in the first well, then serially diluted 1:2 down the plate and incubated overnight at 370C. Plates were washed and then incubated with 50ml anti-mouse IgG1-HRP (1/8000, ab97240, Abcam) for 1 hour at RT and developed with 40ml AEC chromogen substrate kit (AEC101-1KT, Sigma Aldrich). Once wells were sufficiently developed, the plates were washed under running water and dried overnight in the dark before images were captured with ImmunoSpot plate reader (Cellular Technology Ltd). Plates were washed between all steps with PBS-Tw20 5 times and dH2O 3 times.

## Data Availability Statement

All data is available upon reasonable request.

## Ethics Approval for Animal Studies Statement

All research complies with the relevant ethical regulation. All mouse experimentation was performed in the United Kingdom with approval from the Babraham Institute Animal Welfare and Ethical Review Body and complied with European Union and UK Home Office legislation (Home Office License P4D4AF812).

## Author Contributions

L.W. designed and performed experiments, analysed data, constructed the figures, and co-wrote the paper. H.A.C. designed and performed experiments, analysed data, constructed the figures and co-wrote the paper, E.M.W. and S.W. designed and performed experiments. S.E.B. and M.T. provided mice and provided critical feedback. M.A.L. designed experiments and provided critical feedback. All authors contributed intellectually to the work, and reviewed, edited, and approved the paper.

## Acknowledgments

We thank the Babraham Institute Biological Support Unit staff, who performed *in vivo* treatments and animal husbandry. We thank the staff of the Babraham Flow Cytometry and Imaging Facilities for their technical support. This study was supported by funding from the Biotechnology and Biological Sciences Research Council (BBS/E/B/000C0427, Campus Capability Core Grant to the Babraham Institute), a UKRI Frontier award (EP/X022382/1) and a Lister Institute Prize Fellowship to M.A.L,

