## Supplementary Tables for "Expression and role of the RNA-binding protein, ZFP36L1, in mouse T follicular helper cell differentiation and function"

**Supplementary Table 1: Flow Cytometry surface antibodies**

| Antigen | Fluorochrome | Clone | Vendor | Catalogue no. |
|---|---|---|---|---|
| CD90.2 | BUV395 | 53-2.1 | BD Bioscience | 565257 |
| CD19 | BUV661 | 1D3 | BD Bioscience | 612971 |
| CD8a | BUV805 | 53-6.7 | BD Bioscience | 612898 |
| CD44 | BV510 | IM7 | BioLegend | 103044 |
| CD44 | BV711 | IM7 | BioLegend | 103057 |
| CD86 | BV605 | GL-1 | BioLegend | 105037 |
| CD86 | BV510 | GL-1 | BD Bioscience | 563077 |
| CXCR3 | BV650 | 173 | BioLegend | 126531 |
| CD138 | BV711 | 281-2 | BioLegend | 142519 |
| CXCR5 | BV785 | L138D7 | BioLegend | 145523 |
| CD4 | PE-Fire640 | GK1.5 | BioLegend | 100481 |
| PD1 | PE-Cy7 | 29F.1A12 | BioLegend | 135215 |
| CXCR4 | APC | I.276F12 | BioLegend | 146508 |
| IgD | SparkNIR685 | 11-26c-2a | BioLegend | 405750 |
| LiveDead | e780 | - | Invitrogen | 65-0865-14 |
| B220 | APCFire810 | RA3-6B2 | BioLegend | 103278 |

**Supplementary Table 2: Flow Cytometry intracellular antibodies**

| Antigen | Fluorochrome | Clone | Vendor | Catalogue no. |
|---|---|---|---|---|
| IRF4 | Pacific Blue | 3 E4 | BioLegend | 646418 |
| FoxP3 | AF488 | FJK-16s | Invitrogen | 53-5773-82 |
| Tbet | PE-Dazzle594 | 4B10 | BioLegend | 644828 |
| Bcl6 | AF647 | A8 | BioLegend | 648306 |
| Ki67 | AF700 | 16A8 | BioLegend | 652419 |

**Supplementary Table 3: Confocal Microscopy antibodies**

| Antigen | Fluorochrome | Clone | Vendor | Catalogue no. |
|---|---|---|---|---|
| Ki67 | ef450 | SolA15 | Invitrogen | 48-5698-82 |
| PD1 | AF647 | RMP1-30 | BioLegend | 109117 |
| CD3 | AF700 | 17A2 | BioLegend | 100216 |
| CD35 | biotin | 8C12 | BD Biosciences | 553816 |
| IgD | AF700 | 11-26c.2a | BioLegend | 405730 |
| Streptavidin | AF750 | 3 E4 | Invitrogen | S21384 |
